# Profiling copy number variation and disease associations from 50,726 DiscovEHR Study exomes

**DOI:** 10.1101/119461

**Authors:** Evan K. Maxwell, Jonathan S. Packer, Colm O’Dushlaine, Shane E. McCarthy, Abby Hare-Harris, Jeffrey Staples, Claudia Gonzaga-Jauregui, Samantha N. Fetterolf, W. Andrew Faucett, Joseph B. Leader, Andres Moreno-De-Luca, Giusy Della Gatta, Margaret Scollan, Trikaldarshi Persaud, John Penn, Alicia Hawes, Xiaodong Bai, Sarah Wolf, Alexander E. Lopez, Rick Ulloa, Christopher Sprangel, Rostislav Chernomorsky, Ingrid B. Borecki, Frederick E. Dewey, Aris N. Economides, John D. Overton, H. Lester Kirchner, Michael F. Murray, Marylyn D. Ritchie, David J. Carey, David H. Ledbetter, George D. Yancopoulos, Alan R. Shuldiner, Aris Baras, Omri Gottesman, Lukas Habegger, Christa Lese Martin, Jeffrey G. Reid

**Affiliations:** Regeneron Genetics Center, Regeneron Pharmaceuticals, Tarrytown, NY, 10591; Geisinger Health System, Danville, PA, 17822

## Abstract

Copy number variants (CNVs) are a substantial source of genomic variation and contribute to a wide range of human disorders. Gene-disrupting exonic CNVs have important clinical implications as they can underlie variability in disease presentation and susceptibility. The relationship between exonic CNVs and clinical traits has not been broadly explored at the population level, primarily due to technical challenges. We surveyed common and rare CNVs in the exome sequences of 50,726 adult DiscovEHR study participants with linked electronic health records (EHRs). We evaluated the diagnostic yield and clinical expressivity of known pathogenic CNVs, and performed tests of association with EHR-derived serum lipids, thereby evaluating the relationship between CNVs and complex traits and phenotypes in an unbiased, real-world clinical context. We identified CNVs from megabase to exon-level resolution, demonstrating reliable, high-throughput detection of clinically relevant exonic CNVs. In doing so, we created a catalog of high-confidence common and rare CNVs and refined population frequency estimates of known and novel gene-disrupting CNVs. Our survey among an unselected clinical population provides further evidence that neuropathy-associated duplications and deletions in 17p12 have similar population prevalence but are clinically under-diagnosed. Similarly, adults who harbor 22q11.2 deletions frequently had EHR documentation of neurodevelopmental/neuropsychiatric disorders and congenital anomalies, but not a formal genetic diagnosis (i.e., deletion). In an exome-wide association study of lipid levels, we identified a novel five-exon duplication within *LDLR* segregating in a large kindred with features of familial hypercholesterolemia. Exonic CNVs provide new opportunities to understand and diagnose human disease.

## Introduction

Copy-number variants (CNVs), a type of structural variation, are regions of the genome that deviate from the expected normal diploid state as a result of deletion or duplication events. CNVs have been shown to cause or increase risk of a number of diseases^1^. Thus a comprehensive view of CNVs may further our understanding of the genetic bases of human health and disease in service of precision medicine. CNV detection in large clinical and research cohorts is generally performed using array-based technologies given considerations of cost, reliability, and throughput^2^. However, commercially available array-based technologies, or custom-designed arrays used in some clinical laboratories, are limited by probe design and density, achieving a maximum resolution of 5 kb or larger, with calls spanning 50-250 kb being the most reliable. This has additional technical biases and leaves a major class of smaller, exonic CNVs undetected by clinical genomic testing.

While genomic sequencing (whole-exome and whole-genome) is becoming a common diagnostic tool, CNV ascertainment from sequence data is not standard practice due to complexities in reliable detection. Whole-genome sequencing can achieve base-pair resolution^3,4^, but the cost and analytical challenges have thus far limited its application. Exome sequencing provides a balance of resolution, throughput, and cost, making it a more cost-effective choice for clinical labs and population-scale cohort studies, however exome-based CNV detection methods have faced significant technical challenges limiting their adoption^5,6^. We recently developed a high-throughput exome-CNV detection method CLAMMS^7^ that enables reliable common and rare CNV detection across the size spectrum at population-scale. In this study, we sought to explore the phenotypic consequences of exonic CNVs identified in a large clinical care population.

## RESULTS

### Spectrum of CNVs in DiscovEHR Participants

We applied CLAMMS^7^ to the sequenced exomes of 50,726 adult patient-participants who receive health care through the Geisinger Health System (GHS), all of whom consented to participate in the MyCode® Community Health Initiative biobank^8^ and contributed DNA samples for genomic sequencing and analysis as part of the Regeneron-GHS DiscovEHR Study^9^. Each exome is linked to de-identified data extracted from the individual’s electronic health record (EHR), with a median of 13.4 years of longitudinal clinical data available. The sequenced cohort is highly homogeneous in terms of European ancestry (˜97%). While the DiscovEHR Study is not targeted towards any particular age or phenotype, the first 50,000 individuals recruited are enriched for adults (mean age = 61 years) with chronic health problems who interact frequently with the healthcare system, as well as patient-participants sequenced from the Coronary Catheterization Laboratory and the Bariatric Surgery Clinic.

Following application of quality control filters (**Methods**), our high-confidence CNV call set includes 47,349 participants (>93%) and 475,664 CNV events at 13,782 loci (**Supplementary Table S2**). The median size of observed common CNV loci (allele frequency (AF) >= 1%) is 7.1 kb (deletions 4.4 kb, duplications 13.4 kb), compared to 17.8 kb (deletions 8.4 kb, duplications 32.8 kb) in rare CNV loci (AF < 1%). On average, we observed ∼10 high-confidence exonic CNV alleles affecting 14 genes per individual, most of which are common (6.6 deletions and 1.7 duplications, considering only highly mappable regions of the exome). We found an average of one very rare CNV allele per individual (AF < 0.1%; 0.6 duplications and 0.4 deletions; **Fig. 1B, Supplementary Table S2**), and roughly one out of seven individuals possesses at least one CNV that is unique to their exome, relative to the cohort. The vast majority (91%) of CNV loci are extremely rare, observed in fewer than 10 individuals (AF < 0.01%) in our sequenced cohort, with over half representing singletons (**Supplementary Fig. S2**). In total, 13,170 genes are intersected by at least one CNV of less than 2 Mb in length (∼73% of the total callable gene set; **Supplementary Table S2**), and, concordant with other surveys^3,10^, loss-of-function intolerant genes are heavily depleted for observed deletions compared to duplications (**Fig. 1C**, **Supplementary Methods**). In our accompanying linkage disequilibrium map, only half of common CNVs and 30 rare CNVs (0.2%) are well tagged (r^2^ >= 0.3) by a SNP marker, suggesting these CNVs cannot be captured or imputed from genotyping arrays (**Supplementary Methods**).

**Figure 1:**
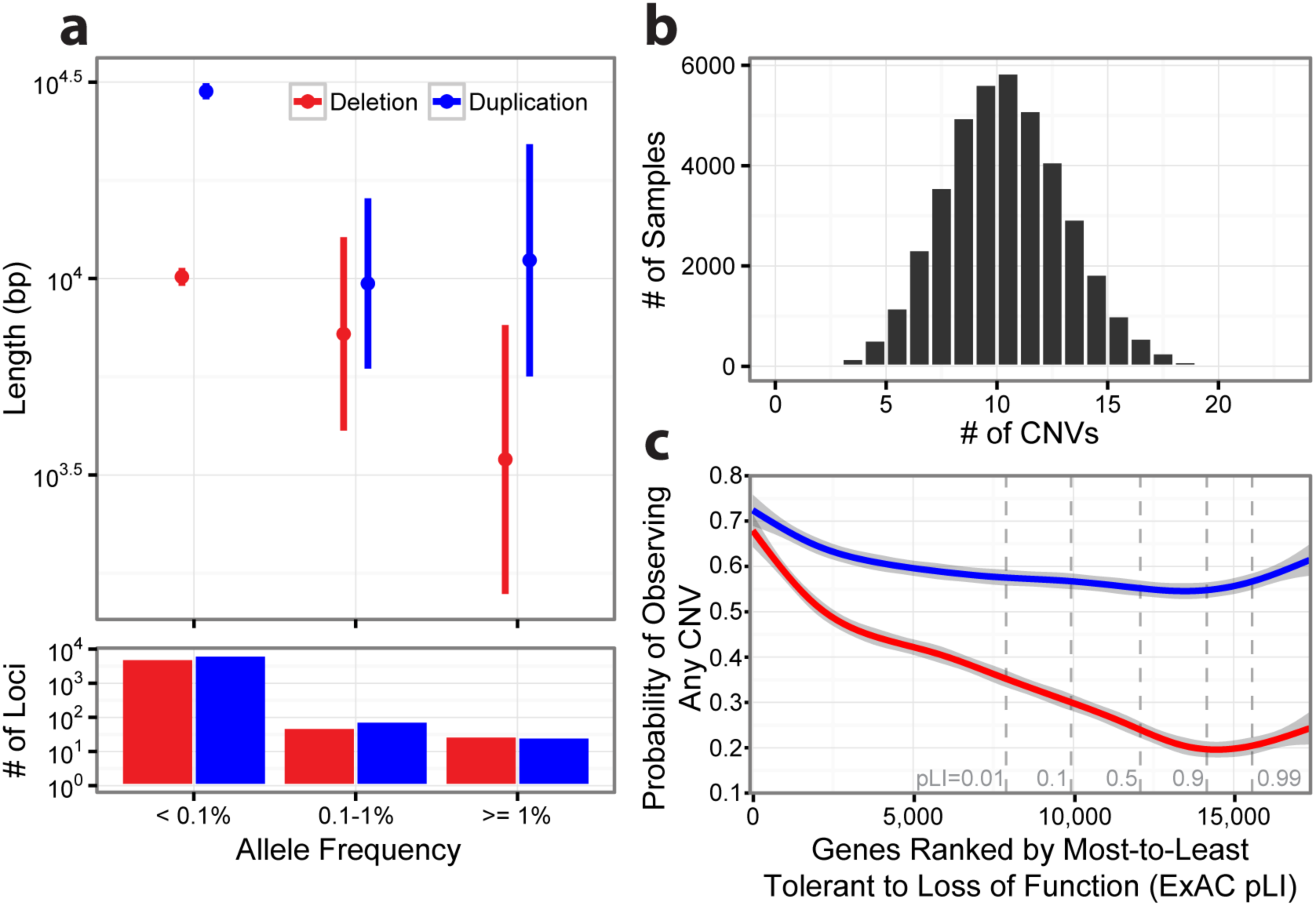
Exome-wide survey of high-confidence CNVs from 47,349 individuals. **A)** Mean length (95% confidence bands) for deletion and duplication loci at varying allele frequency ranges. **B)** Sample-wise distribution of CNV count (**Supplementary Table S2**). **C)** Comparison of CNV intolerance in genes relative to loss-of-function (LOF) intolerance probabilities computed from single nucleotide variants (ExAC pLI metric v0.3^30^). While duplications are observed consistently in ∼60% of genes regardless of LOF intolerance, observed deletion frequencies decrease in concordance with LOF intolerance.

### Segregation of CNVs in Computationally Inferred Kindreds

Cross-referencing CNVs with computationally inferred pedigrees (**Supplementary Methods**) enables tracking of familial variants as well as analysis of transmission rates, providing estimates of cohort-wide call sensitivity and specificity. The call set exhibits an average parent-child transmission rate of ∼46.5% for rare heterozygous CNVs, consistent with a theoretical rate of 50% assuming equal probability of transmission and lack of transmission disequilibrium. This includes many rare small CNVs (<= 3 exons, 0.54 per sample cohort-wide) transmitted at 42.17%. We validated, using qPCR, a sample of 100 high-confidence CNVs identified in 259 parent-child duos (**Methods**), estimating precision and recall to be 98% and 92%, respectively, in the smallest class of exonic CNVs (1-3 exons). Comparatively, an alternative (and widely used) exome CNV calling method, XHMM^11^, achieved only 36.1% recall (1% for single-exon calls) with 96.8% precision in this size range (**Supplementary Table S3 & Methods**).

### Prevalence and Clinical Impact of Disease-associated Exonic CNVs

We surveyed frequencies of a selected set of known disease-associated CNVs likely reflecting the spectrum of coding CNVs expected in a broad, predominately European hospital-based population (**Table 1**). This EHR-linked resource provided the opportunity to evaluate penetrance and expressivity of Mendelian disease-associated CNVs and to characterize the clinical consequences of other exonic CNVs among a broadly sampled adult population.

**Table 1:**
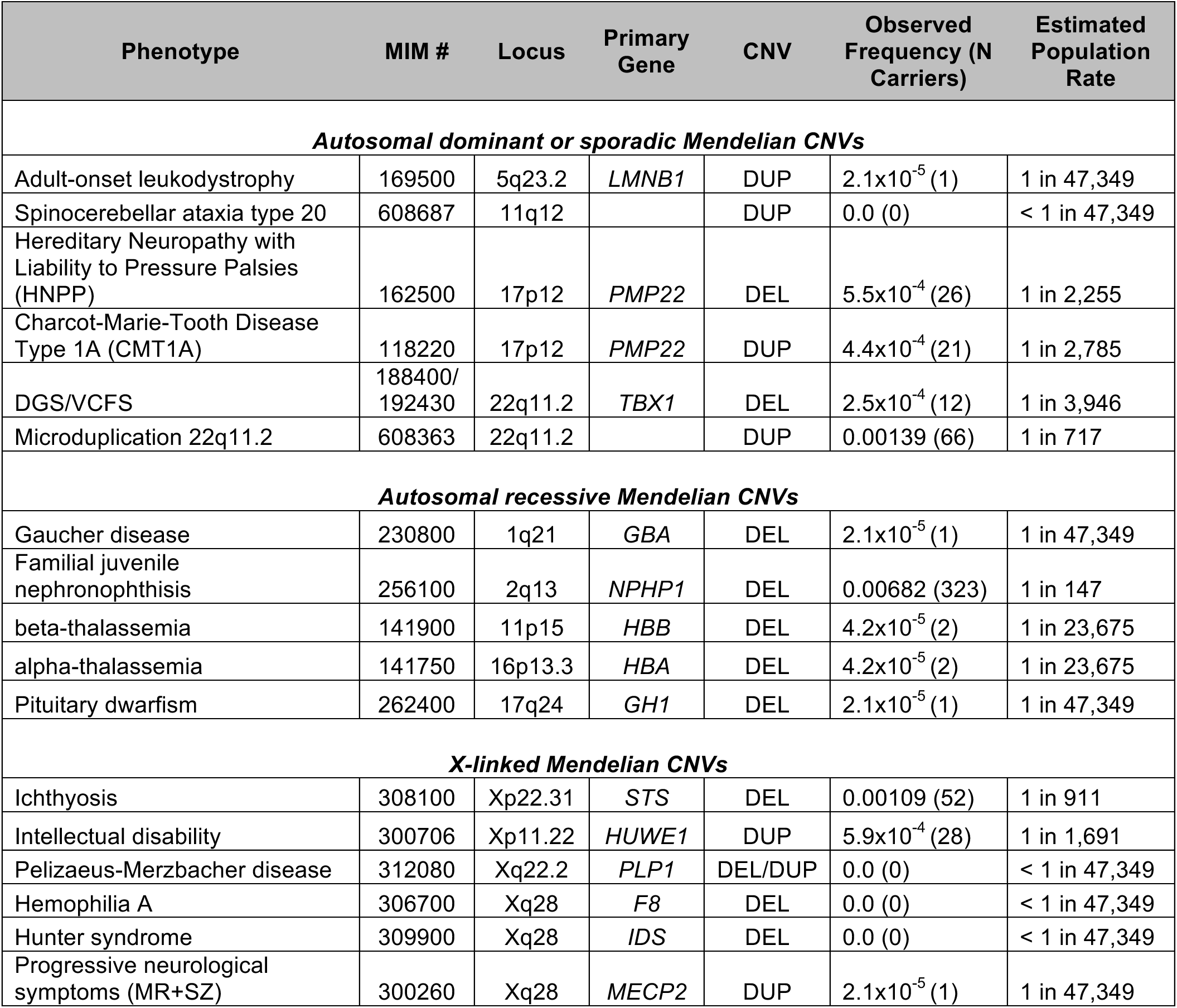
Observed frequencies of select known disease-associated CNV loci.

| Phenotype | MIM # | Locus | Primary Gene | CNV | Observed Frequency (N Carriers) | Estimated Population Rate |
| --- | --- | --- | --- | --- | --- | --- |
| ***Autosomal dominant or sporadic Mendelian CNVs*** |  |  |  |  |  |  |
| Adult-onset leukodystrophy | 169500 | 5q23.2 | *LMNB1* | DUP | $2.1 \times 10^{-5}$ (1) | 1 in 47,349 |
| Spinocerebellar ataxia type 20 | 608687 | 11q12 |  | DUP | 0.0 (0) | < 1 in 47,349 |
| Hereditary Neuropathy with Liability to Pressure Palsies (HNPP) | 162500 | 17p12 | *PMP22* | DEL | $5.5 \times 10^{-4}$ (26) | 1 in 2,255 |
| Charcot-Marie-Tooth Disease Type 1A (CMT1A) | 118220 | 17p12 | *PMP22* | DUP | $4.4 \times 10^{-4}$ (21) | 1 in 2,785 |
| DGS/VCFS | 188400/<br>192430 | 22q11.2 | *TBX1* | DEL | $2.5 \times 10^{-4}$ (12) | 1 in 3,946 |
| Microduplication 22q11.2 | 608363 | 22q11.2 |  | DUP | 0.00139 (66) | 1 in 717 |
| ***Autosomal recessive Mendelian CNVs*** |  |  |  |  |  |  |
| Gaucher disease | 230800 | 1q21 | *GBA* | DEL | $2.1 \times 10^{-5}$ (1) | 1 in 47,349 |
| Familial juvenile nephronophthisis | 256100 | 2q13 | *NPHP1* | DEL | 0.00682 (323) | 1 in 147 |
| beta-thalassemia | 141900 | 11p15 | *HBB* | DEL | $4.2 \times 10^{-5}$ (2) | 1 in 23,675 |
| alpha-thalassemia | 141750 | 16p13.3 | *HBA* | DEL | $4.2 \times 10^{-5}$ (2) | 1 in 23,675 |
| Pituitary dwarfism | 262400 | 17q24 | *GH1* | DEL | $2.1 \times 10^{-5}$ (1) | 1 in 47,349 |
| ***X-linked Mendelian CNVs*** |  |  |  |  |  |  |
| Ichthyosis | 308100 | Xp22.31 | *STS* | DEL | 0.00109 (52) | 1 in 911 |
| Intellectual disability | 300706 | Xp11.22 | *HUWE1* | DUP | $5.9 \times 10^{-4}$ (28) | 1 in 1,691 |
| Pelizaeus-Merzbacher disease | 312080 | Xq22.2 | *PLP1* | DEL/DUP | 0.0 (0) | < 1 in 47,349 |
| Hemophilia A | 306700 | Xq28 | *F8* | DEL | 0.0 (0) | < 1 in 47,349 |
| Hunter syndrome | 309900 | Xq28 | *IDS* | DEL | 0.0 (0) | < 1 in 47,349 |
| Progressive neurological symptoms (MR+SZ) | 300260 | Xq28 | *MECP2* | DUP | $2.1 \times 10^{-5}$ (1) | 1 in 47,349 |

We identified 12 patient-participants with 22q11.2 deletion syndrome (MIM #188400), two of which are from the same family. This deletion is one of the most common and clinically documented recurrent CNVs; our observed prevalence of 1 in 3,946 in the DiscovEHR cohort is in line with previous estimates of 1 in 4,000 live births^12^. Of the 12 participants with a 22q11.2 deletion, 10 (83%) had ICD-9 codes consistent with the 22q11.2 deletion syndrome phenotype, including 8 (67%) with a diverse set of neurodevelopmental/neuropsychiatric disorders (NDD) and 6 (50%) with congenital anomalies, with 5/12 (42%) participants having both NDD and congenital anomaly phenotypes reported (**Supplementary Methods).** However, only one patient-participant (age 50 years old) with a 22q11.2 deletion had a specific genetic diagnosis of velocardiofacial syndrome (ICD 758.32) documented in their EHR, suggesting that most individuals are likely unaware of the genetic etiology of their clinical phenotypes.

We also found 25 patient-participants with 17p12 duplications that include the *PMP22* gene; 21 having duplications spanning the common recurrent CNV locus associated with the most common form of peripheral neuropathy, Charcot-Marie-Tooth disease type 1A (CMT1A; MIM #118220)^13^, three with atypical duplications in two families, and one with breakpoints that could not be reliably estimated (**Supplementary Fig. S12**). Fourteen out of 21 of the patient-participants with a common recurrent duplication and 2/3 with an atypical duplication (a parent-child duo) had a clinical diagnosis of CMT1A. Seven out of 21 individuals with a recurrent duplication had non-specific diagnoses consistent with CMT1A recorded in their EHR (**Supplementary Methods**). Three patient-participants had neuropathy phenotypes attributed to type-2 diabetes, suggesting a possible misdiagnosis of the primary hereditary neuropathy. Similarly, we identified and validated 26 patient-participants with the reciprocal 17p12 deletion associated with Hereditary Neuropathy with liability to Pressure Palsies (HNPP; MIM #162500)^13^, all of which spanned the common recurrent CNV region. However, only 1/26 patient-participants had a clinical diagnosis of HNPP in their EHR and 6 had consistent, but non-specific diagnoses. Our observed population frequencies for the recurrent CMT-associated duplication (1/2,255 total, 1/2,785 *de novo;* **Supplementary Methods**) and the related HNPP-associated deletion (1/1,821 total, 1/2,255 *de novo*) are roughly equal, but their EHR-based diagnostic rates are vastly different, suggesting that the deletion is largely clinically under-diagnosed. Forty-nine patient-participants with 17p12 CNVs and five non-carrier relatives were validated with qPCR, demonstrating accurate identification of this clinically relevant CNV from the exome data (**Supplementary Fig. S12 & Methods**).

Finally, among 323 patient-participants with 2q13 deletions encompassing the entirety of the nephronophthisis-1 (*NPHP1*) gene, we identified two individuals with homozygous deletions. Both of these individuals (age 31 and 38 years) had a documented diagnosis of medullary cystic kidney (ICD 753.16) and end stage renal disease (ICD 585.6) requiring transplantation, consistent with autosomal recessive juvenile nephronophthisis (OMIM #256100). These results exemplify the importance of differentiating heterozygous versus homozygous deletions for clinical correlations.

### Exome-wide analysis of associations of exonic CNVs with serum lipid levels

We next tested the ability to identify novel disease susceptibility loci through the intersection of exome CNVs and EHR-derived phenotypes. We performed an exome-wide association study of our CNV loci with fasting serum lipid levels (**Methods**), which are heritable risk factors for ischemic cardiovascular disease, in a subset of 39,087 individuals with available lipid data. At a Bonferroni-corrected significance threshold of p<1.2×10^-5^, three CNV loci significantly associated with lipid levels (**Table 2**). We then evaluated the penetrance of these lipid-associated variants to coronary artery disease (CAD).

**Table 2:**
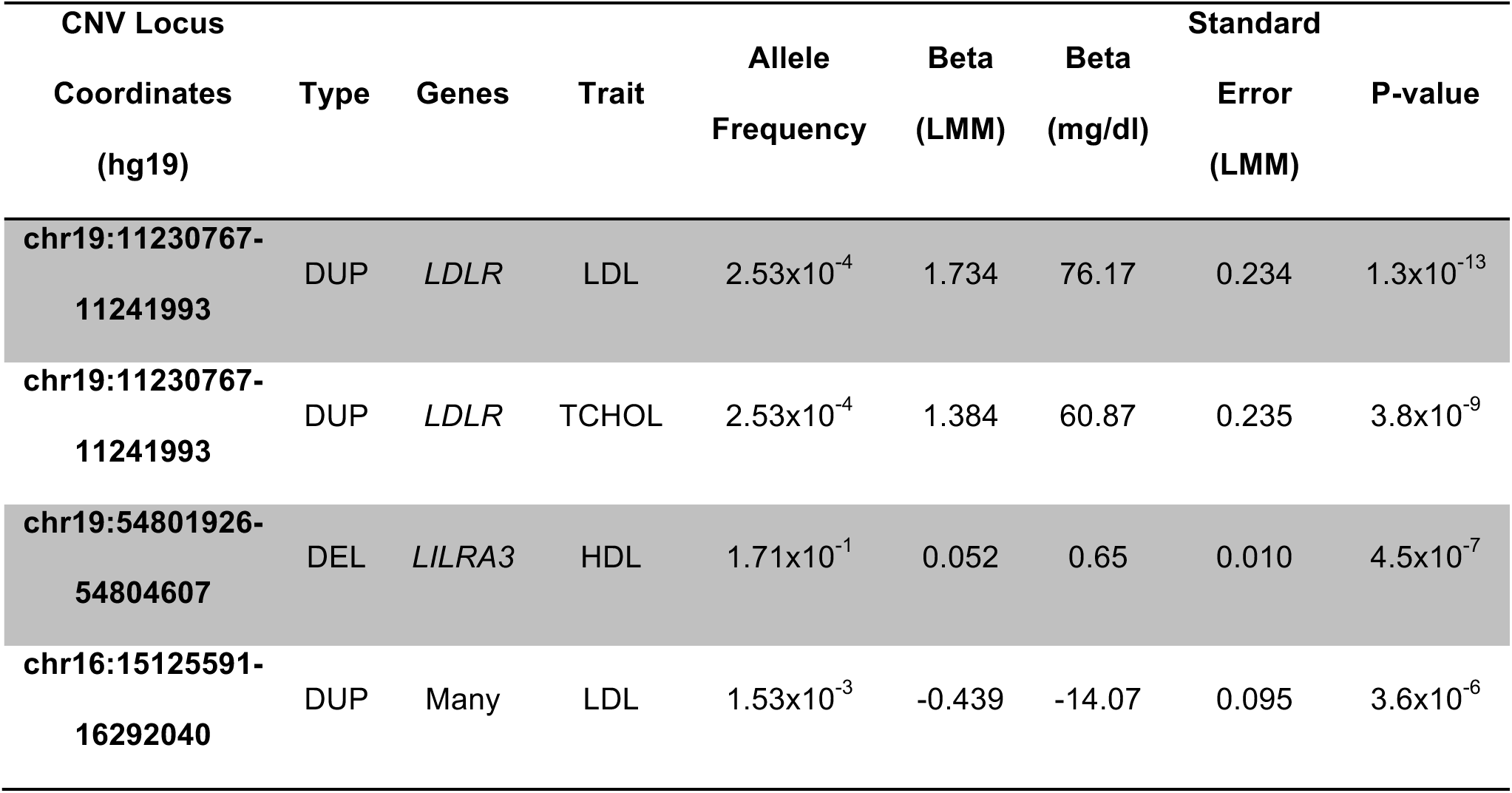
**Exome-wide significant associations between high-confidence exonic CNV loci and EHR-derived serum lipid traits (LDL-c, HDL-c, total cholesterol “TCHOL”, and triglycerides).** LMM = Linear Mixed Model (**Methods**).

A novel duplication of g.chr19:11230767-11241993 (hg19/GRCh37) encompassing exons 13-17 of the low-density lipoprotein receptor gene *LDLR* was identified in 29 individuals and found to be associated with high LDL cholesterol (*β*=1.73 [76 mg/dl], p=1.3×10^-13^) and high total cholesterol (β=1.38 [61 mg/dl], p=3.8×10^-9^; **Methods**). Breakpoints were identified in *Alu* repeats within introns 12 and 17 using whole-genome sequencing, confirming a tandem insertion site predicted to occur in-frame (**Fig. 2A**). The duplication was validated in all carriers by Sanger sequencing (**Supplementary Fig. S6**). CNVs have previously been reported in LDLR^14^; however, this duplication appears to be novel. In overexpression experiments in HEK293 cells (**Supplementary Methods**), *LDLR* containing the exon 13-17 tandem duplication (Dup13-17) produced a larger protein product than the wild-type (WT) receptor that accumulates in the endoplasmic reticulum. As a result, the mutant receptor is not expressed on the cellular surface and has no LDL uptake activity. Thus, the duplication behaves similarly to the known FH-causing p.G549D point mutation, classified as a transport-inhibiting (class-2) mutation (**Figs. 2B-E & S13**). Taken together, this evidence suggests that the duplication causes loss-of-function of LDL receptor activity.

**Figure 2:**
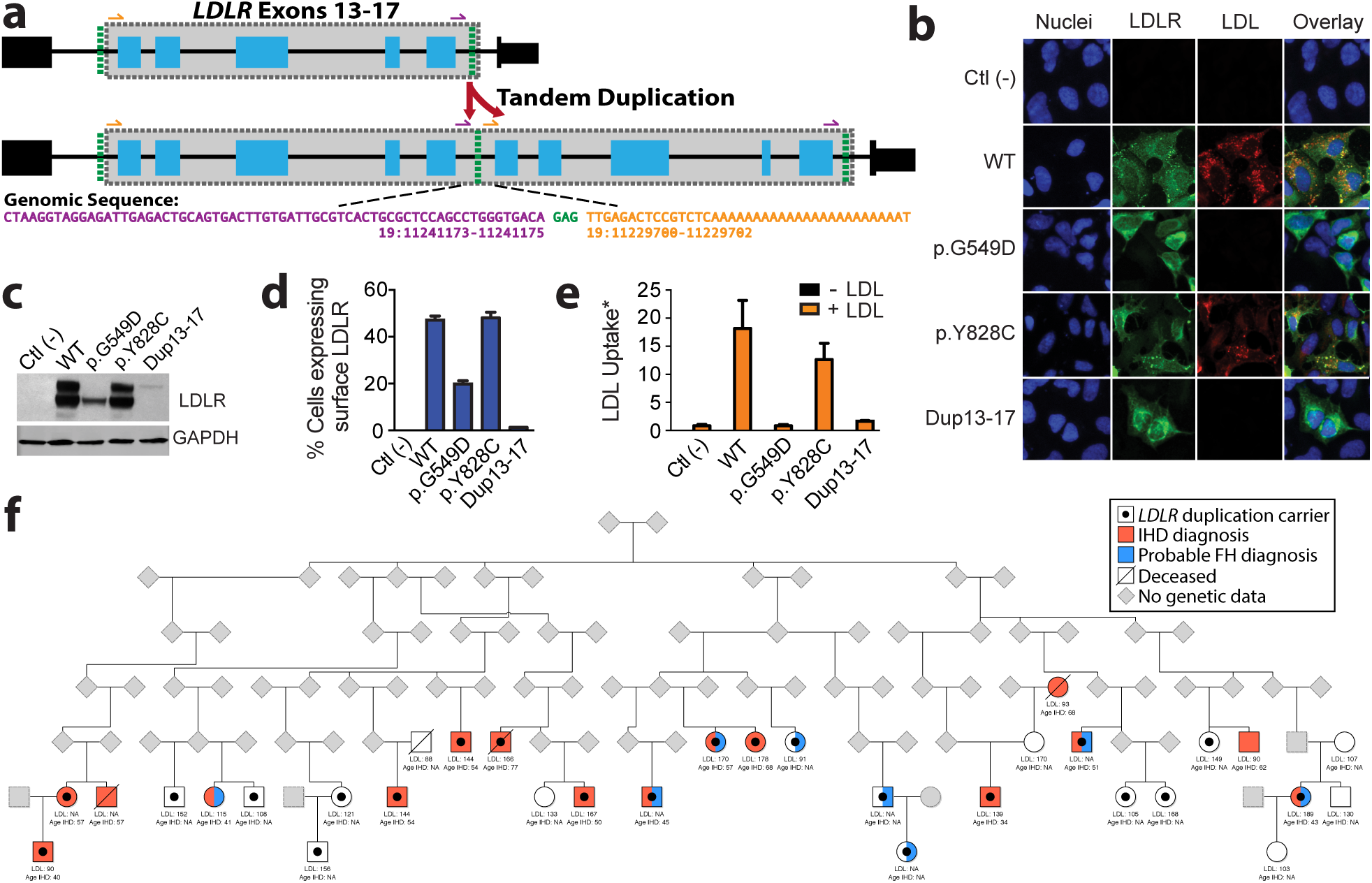
**A familial hypercholesterolemia-associated tandem duplication within *LDLR* causes loss-of-function and segregates with high LDL cholesterol and heart disease A)** Whole-genome sequencing and Sanger validation (**Supplementary Fig. S6**) confirmed the *LDLR* duplication in 29 individuals occurs in tandem encompassing exons 13-17 (“Dup13-17”). Both the breakpoint and insertion loci occur in intronic *Alu* repeat sequences, having a shared three nucleotide microhomology (green). The predicted protein translation is in-frame, creating a larger mutant receptor (**C**) that is not expressed on the cellular surface (**B & D, Supplementary Fig. S13**), resulting in significantly reduced LDL uptake activity and loss-of-function (**B & E**). **F)** Reconstructed pedigree estimate containing 22/29 carriers of the duplication and ten unaffected related (first or second degree) individuals from the sequenced cohort. Five additional carriers (not drawn) are included in this pedigree (**Supplementary Methods**). Elevated LDL and total cholesterol as well as increased prevalence of coronary artery disease and early-onset ischemic heart disease (“Age IHD” <55 for males and <65 for females) segregate with duplication carriers; but only eight carriers have a probable FH diagnosis coded in their EHRs (**Supplementary Methods**). *LDL Uptake = Cytoplasmic LDL Puncti / # GFP+ Cells.

The *LDLR* duplication associates with markedly increased CAD risk (OR=5.01, Logistic regression, p=6×10^-4^). Using pedigree reconstruction and distant-relatedness analysis (**Supplementary Methods**), we connected 27/29 carriers into a single large estimated pedigree, dating their common ancestor to at least six generations ago and identifying 10 related individuals not harboring the *LDLR* duplication (**Fig. 2C**). In this extended pedigree, the duplication segregated with high LDL levels and 15/29 duplication carriers had ischemic heart disease (IHD) diagnoses, 11 of who presented with early-onset IHD (**Supplementary Methods**). Given the genetic and functional evidence, and the observation that familial hypercholesterolemia (FH) cases are frequently attributed to *LDLR* mutations^14^, we conclude that this is a novel pathogenic FH-causing CNV. Present in 1/1,749 samples, this novel *LDLR* duplication represents roughly 13% of the pathogenic FH variants (30% of those specific to *LDLR*) observed in our cohort, which have an overall prevalence of 1:256 unselected individuals in sequenced participants^15^. Notably, only eight patient-participants carrying the *LDLR* duplication and one relative without the duplication have a probable FH diagnosis coded in their EHR (**Supplementary Methods**).

We also identified a common deletion encompassing the last 6/7 exons of the leukocyte immunoglobulin (Ig)-like receptor A3 gene (*LILRA3;* AF ≅ 17%, consistent with previous estimates in Europeans^16^), that was associated with increased HDL levels (*β*=0.05 [0.65 mg/dl], p=4.5x10^-7^, **Methods**). No association with the prevalence of CAD was observed. While the size of this variant (∼2.6 kb exonic, 6.7 kb genomic) is not amenable to detection with standard clinical CNV arrays, CLAMMS calls at this locus were previously qPCR-validated in 165 samples (69 CNV carriers) with perfect sensitivity and specificity^7^. Genome-wide association studies (GWAS) have identified SNVs adjacent to *LILRA3* that associate with HDL levels^17^; our analysis revealed that the deletion and GWAS SNV (rs386000) are in linkage disequilibrium (r^2^=0.77, D’=0.959; **Supplementary Dataset S2**). Multiple expression quantitative trait loci (eQTLs) have been mapped to *LILRA3*, but the deletion is likely to be driving the effect^18^ and contributing to the variation in HDL levels. This deletion has previously been investigated for association with other diseases^16^; we observed nominally increased risk for rheumatoid arthritis (p=0.041, OR=1.1,95% confidence interval=[1.004,1.223], logistic regression) and decreased risk for prostate malignancies (p=0.045, OR=0.9, 95% confidence interval=[0.797,0.997], logistic regression) among carriers (**Supplementary Methods**). Further testing of this *LILRA3* CNV in other well-powered cohorts will shed light into the true genotype-phenotype associations. In addition to *LILRA3* deletions, we demonstrate the ability to detect a small, complex CNV in the haptoglobin gene (*HP*) through a single mappable exon well enough to recapitulate the directionality of previously observed associations with increased LDL and total cholesterol^19^ (**Supplementary Methods**).

## DISCUSSION

Our study demonstrates the utility of exome sequencing data for analyzing common and rare CNVs in the emerging paradigm of precision medicine using EHR-derived health information. We show robust detection of small exonic CNVs well below the size threshold of array-based technologies (**Supplementary Methods**), providing further evidence that the density of markers on genotyping arrays is insufficient for characterizing the full CNV spectrum in humans and necessitates higher resolution methods (**Supplementary Fig. S4**).

Our catalog of exonic CNVs represents a substantial source of genomic variation in the DiscovEHR study population and exemplifies the clinical value of CNVs. We survey the clinical expressivity of known pathogenic CNVs, and via exome-wide associations with serum lipid traits, discover a novel duplication in *LDLR* causing familial hypercholesterolemia (FH). Among both known and novel pathogenic CNVs, the genetic basis for disease is frequently under-diagnosed. Clinical variability is common among individuals with known recurrent CNVs (e.g. 17p12 CNVs and 22q11.2 deletion syndrome), presenting a diagnostic challenge to clinicians, especially in patients with more mild symptoms and those born before the widespread use of clinical genetic testing. In the case of the *LDLR* duplication, this novel CNV is undetectable with standard genetic screening platforms; thus, the handful of carriers with FH-consistent EHR documentation were diagnosed based on clinical features, not the underlying genetic cause. Combined with other reports of *LDLR* rearrangements^14,20^ and the density of *Alu* repeat elements (2.24 × 10^-3^ per base intronic sequence, 99.4^th^ percentile for intron-containing genes; **Supplementary Methods**), *LDLR* may be particularly prone to genomic rearrangements^21^. These data highlight the potential for high-throughput CNV screening to improve diagnostic rates for FH and inform patient treatment.

With the growing prevalence of whole-exome and whole-genome sequencing, the substantial body of literature implicating CNVs in both rare and common disease, and the magnitude of copy-number variation we observe in the exome, the inclusion of methods to identify CNVs and other structural variants within standard sequence-based informatics pipelines is long overdue. In addition, the ability to identify rare homozygous deletions (e.g. 2q13) in large cohort studies, like DiscovEHR, will contribute to human gene knockout catalogs and the identification of autosomal recessive conditions. As we find that over 90% of distinct CNVs are present in less than 1/5,000 individuals; large sample sizes or targeted recruitment of additional family members are essential to establish phenotypic associations. Furthermore, detection of exonic and larger clinically relevant CNVs from exome sequence data will streamline genetic testing so that both sequence and copy number variants can be called from one test methodology. The ability to obtain a more comprehensive view of human genetic variation, both per individual and across populations, will facilitate the advent of precision medicine.

## METHODS

### Study population

The human genetics studies were conducted as part of the DiscovEHR study of the Regeneron Genetics Center and the Geisinger Health System. Patient-participants who receive health care through Geisinger Health System were consented to participate in the MyCode Community Health Initiative and DiscovEHR cohort following an IRB approved protocol^8^. DNA samples and exome data from 50,726 adult patient-participants were included in this study. Detailed information on the clinical characteristics of this cohort can be found in^22^. The Regeneron Genetics Center funded study sample collection, sequence data generation, and clinical and sequence data analysis. All participants gave informed written consent.

### Sample preparation and sequencing

Genomic DNA samples were transferred to the Regeneron Genetics Center from the Geisinger Health System in 2D matrix tubes (Thermo Scientific, Waltham, MA), logged into our LIMS (Sapio Sciences, Baltimore, MD), and stored in our automated biobank at -80°C (LiCONiC TubeStore, Woburn, MA). Sample quantity was determined by fluorescence (Life Technologies, Carlsbad, CA) and quality assessed by running 100ng of sample on a 2% pre-cast agarose gel (Life Technologies). The DNA samples were normalized and one aliquot was sent for genotyping (Illumina Inc., San Diego, CA, Human OmniExpress Exome Beadchip) and another sheared to an average fragment length of 150 base pairs using focused acoustic energy (Covaris LE220, Woburn, MA). The sheared genomic DNA was prepared for exome capture with a custom reagent kit from Kapa Biosystems (Wilmington, MA) using a fully-automated approach developed at the Regeneron Genetics Center (Tarrytown, NY). A unique 6 base pair barcode was added to each DNA fragment during library preparation to facilitate multiplexed exome capture and sequencing. Equal amounts of sample were pooled prior to exome capture with NimbleGen (Roche NimbleGen, Madison, WI) probes (SeqCap VCRome). Captured fragments were bound to streptavidin-conjugated beads and non-specific DNA fragments removed by a series of stringent washes according to the manufacturer’s recommended protocol (Roche NimbleGen). The captured DNA was PCR amplified and quantified by qRT-PCR (Kapa Biosystems). The multiplexed samples were sequenced using 75 bp paired-end sequencing on an Illumina v4 HiSeq 2500 to a coverage depth sufficient to provide greater than 20x haploid read depth of over 85% of targeted bases in 96% of samples (approximately 80x mean haploid read depth of targeted bases).

Upon completion of sequencing, raw data from each Illumina Hiseq 2500 run was gathered in local buffer storage and uploaded to the DNAnexus (Mountain View, CA) platform^23^ for automated analysis. Sample-level read files were generated with CASAVA (Illumina Inc.) and aligned to GRCh37.p13 with BWA-mem^24,25^. The resultant BAM files were processed using GATK^26^ and Picard to sort, mark duplicates, and perform local realignment of reads around putative indels.

### CNV calling and quality control

CLAMMS^7^ was applied to the exomes of all samples in parallel within a distributed compute environment. CLAMMS calls CNVs from each sample’s source BAM file using 1) base-level depth-of-coverage calculations from aligned reads having mapping quality >= 30, and 2) seven sequencing quality control (QC) metrics computed using Picard (http://broadinstitute.github.io/picard/). Depth-of-coverage profiles are generated for exon “windows” representing either the entire exon or a 500-1000 base pair contiguous segment of an exon for long exons. Exon window coverage distributions are normalized independently for every sample, adjusting for GC content and overall sequencing depth. CLAMMS detects CNVs by comparing a sample’s normalized coverage profile to a reference panel of 100 samples matched with respect to QC metrics via a k-nearest neighbors search, where expected coverage distributions are modelled with exon-window specific mixture models linked together with a genomic distance-aware Hidden Markov Model (HMM).

Extensive quality control procedures were applied to the call set with the goal of producing a set of CNV loci exhibiting high specificity (see **Supplementary Methods**). Briefly, the sample set was filtered of “outlier” samples having inflated CNV rates (>28 CNVs, 2x median) resulting in a high-confidence call set containing 47,349 samples (93%). CNVs in the remaining samples were annotated with: 1) two model-based metrics from CLAMMS (Q_non___dip_ and Q_exact_, representing confidence measures that the region is non-diploid and consistent with the exact called copy number, respectively), 2) allele balance and zygosity of SNVs within called CNV regions, 3) the frequency of overlapping CNVs among outlier samples, and 4) the quality and zygosity measures for all non-outlier carriers of the locus (representing cohort-wide locus performance). Filtering criteria based on these annotations were trained with respect to the cohort-wide transmission rate of rare heterozygous CNVs among parent-child duos from genetically reconstructed pedigrees generated with PRIMUS^27^. The training set was targeted to 47.5% transmission, achieving 47.36% transmission in the training set, 46.02% in the test set, and 46.59% combined. This includes several small (1-3 exon) CNVs (∼29% of loci) transmitted at 42.17%, with single-exon CNVs (∼8% of loci) being transmitted at 38.96% (**Supplementary Table S1**).

### CNV validation data

In a prior study^7^, we validated select common and rare CNV loci identified by CLAMMS using TaqMan qPCR and demonstrated higher performance and resolution than alternative exome CNV callers as well as CNV detection from SNP arrays. In the present study, we have expanded our validation set to include additional TaqMan qPCR validation on a representative set of 100 high-confidence CNV loci identified in 259 parent-child duos, with a focus on small CNVs (1-3 exon; 86/100 loci) having at least one duo in which a non-transmission is predicted from CLAMMS (i.e. present in the parent, absent in the child). This provided an unbiased method to test the sensitivity and specificity of CLAMMS at high-confidence small CNV loci, particularly with respect to interpretation of the cohort-wide transmission rate estimates (42.17% for 1-3 exon CNVs, 38.96% for single-exon CNVs; **Supplementary Table S1**). Among the small CNV loci, we observed 10.5% of non-transmission calls did not qPCR validate, corresponding to a 2.2% FPR, 8% FNR, and a total error rate of 5.3% (1-accuracy). Among single-exon CNVs, we observed a 3.5% FPR, 9.2% FNR, and a total error rate of 6.6% (**Supplementary Table S3**).

In addition to TaqMan qPCR validations, a large subset of samples (N=34,246) were processed with Illumina HumanOmniExpressExome-8v1-2 genotyping arrays in twelve batches and subsequently tested with PennCNV^28^. PennCNV calls were quality controlled using a 95^th^ percentile cutoff applied to each batch for samples having extreme Log-R Ratio standard deviation (LRR_SD) and B Allele Frequency (BAF) drift. Calls at neighboring loci were considered for merging using the clean_cnv module with the ‐‐bp combine_seg flag. We tested the transmission rate of PennCNV calls from samples in identified parent-child duo relationships having fewer than 50 CNVs (N=21,792), as excessive numbers of CNVs often indicate low sample quality, high rates of *de novo* mutation, or somatic copy number variation that would confound transmission rate calculations. Consistent with our previous observations^7^, we found that PennCNV calls were only reliable (i.e. produced transmission rates close to 50%) at size thresholds approaching >100 Kb or larger (**Supplementary Figure S4**).

### Statistical Analysis

Associations between CNVs and lipid traits were performed using a linear mixed model implemented in BOLT-LMM^29^ with a genetic relationship matrix included as a random effect. Deletions and duplications at the same locus were considered separately. Median values for serially measured laboratory traits, including total cholesterol, low-density lipoprotein cholesterol (LDL-C), high-density lipoprotein cholesterol (HDL-C) and triglycerides were calculated for all individuals with two or more measurements in the EHR following removal of likely spurious values that were > 3 standard deviations from the intra-individual median value. For the purposes of exome-wide association analysis of serum lipid levels, total cholesterol, LDL-C, triglycerides, and HDL-C were adjusted for lipid-altering medication use by dividing by 0.8, 0.7, 0.82, and 1.044, respectively, to estimate pre-treatment lipid values. Medication-adjusted HDL-C and triglycerides were log_10_ transformed, and medication-adjusted LDL-C and total cholesterol values were not transformed. We then calculated trait residuals after adjustment for age, age^2^, sex, and the first ten principal components of ancestry, and rank-inverse-normal transformed these residuals prior to exome-wide association analysis (**Supplementary Methods**).

## Acknowledgements

The authors would like to thank the MyCode Community Health Initiative participants for their permission to utilize their health and genomics information in the DiscovEHR collaboration and for their ongoing engagement in multiple facets of this research. The authors also thank David D’Ambrosio and Jennifer Espert for performing the TaqMan qPCR validation experiments and Dr. Scott Myers for assisting with clinical correlations. This research was funded by the Regeneron Genetics Center, a wholly-owned subsidiary of Regeneron Pharmaceuticals. This study was also partially supported by grant RO1MH074090 (Drs. Ledbetter and Martin) from the National Institute of Mental Health. Regeneron manufactures and markets Praluent (alirocumab), a *PCSK9* inhibiting antibody, for treatment of familial hypercholesterolemia and other indications. Under the relevant agreement with the Geisinger Health System (GHS), Regeneron is prohibited from making genomic DNA samples and individual level phenotype information obtained from the GHS MyCode-DiscovEHR Project available to another party, and Regeneron is prohibited from disclosing a copy of the agreement to another party. A patent application on the CLAMMS copy number variant calling method has been filed by Regeneron, and a patent application that contains data in the manuscript has been filed by Regeneron.

